# Metformin antiproliferative activity is exclusively mediated by the membrane functional expression of the Chloride Intracellular Channel 1 in glioblastoma stem cells

**DOI:** 10.1101/2022.12.31.522371

**Authors:** Ivan Verduci, Francesca Cianci, Riccardo Cazzoli, Gaetano Cannavale, Marina Veronesi, Beatrice Balboni, Matteo Ranucci, Luca Maria Giovanni Palloni, Cristina Richichi, Stefania Faletti, Daniela Osti, Giorgia Ailuno, Gabriele Caviglioli, Carlotta Tacconi, Stefania Girotto, Federica Barbieri, Alessandro Fantin, Andrea Cavalli, Giuliana Pelicci, Tullio Florio, Saverio Minucci, Michele Mazzanti

## Abstract

Metformin is the first-line drug for type-2 diabetes. Retrospective analyses, based on diabetic patients’ clinical data, demonstrate that daily assumption of metformin reduces the incidence of several kinds of solid tumors. Even though it is widely agreed that metformin must be internalized to accomplish its pharmacological activity, direct evidence about metformin membrane permeability and/or the presence of a specific membrane receptor in cancer cells is still missing. Here, we show that the transmembrane form of Chloride Intracellular Channel 1 (tmCLIC1) works as a privileged metformin receptor in glioblastoma stem-like cells. We found that metformin impairs tmCLIC1 activity by a specific binding coordinated by arginine 29. Its mutation, preventing metformin to bind and block tmCLIC1, abolishes the biguanide inhibition of glioblastoma cell proliferation in 2D and 3D models and metformin dependent effect on mitochondrial respiration. In addition, we demonstrate the direct binding between the drug and its target, and by *in vivo* experiments on zebrafish embryos and mice orthotopically engrafted with glioblastoma cells and treated with metformin, we prove that metformin binding to tmCLIC1 is crucial for metformin antineoplastic effect. Considering tmCLIC1’s contribution to glioblastoma progression, the present work provides the fundaments for future development of strategies aimed at improving metformin-tmCLIC1 interaction to further increase metformin therapeutic potential.

## Introduction

Glioblastoma (GBM, glioma grade 4 WHO^1^), is the deadliest brain tumor, representing a major clinical challenge. Despite the standard of care and cutting-edge strategies^2–6^, patients’ life expectancy stands invariantly poor^7,8^. Tumor heterogeneity and its extensive invasiveness make GB difficult to be radically resected by neurosurgery and recurrence are common. There is scientific consensus that tumor relapse originates from progenitor/stem-like cells known as glioblastoma stem cells (GSCs). GSCs are endowed with distinct tumorigenic potential displaying properties of self-renewal and multi-lineage differentiation that contribute to tumor mass heterogeneity^9–14^. Thus, GSCs represent an appealing target for newly designed therapies.

In recent years, metformin’s anti-cancer properties have been highlighted. Metformin administration to type 2 diabetes (T2D) patients has been positively linked with a decrease in the risk of several cancer developments and cancer-related mortality, including glioblastoma^15–18^. Additionally, numerous studies on glioblastoma demonstrated that metformin anticancer properties are directed against GSCs^19–23^.

Despite this evidence, metformin’s mechanism of action on cancer cells has not been clarified yet, although it has been proposed to target the oxidative phosphorylation (OXPHOS) pathway^24–26^. GSCs are able to shuttle between glycolysis and OXPHOS energetic pathways^27–29^, and this metabolic plasticity can be the major reason for metformin’s low effectiveness in clinical trials so far^30^. However, metformin-treated GSCs show a parallel and not-additive inhibition of proliferation to that obtained by impairing the activity of the transmembrane form of the Chloride Intracellular Channel 1 (tmCLIC1) with pharmacological tools^31,32^.

CLIC1 is a metamorphic protein that is predominantly cytoplasmic under physiological conditions, while, in response to persistent stress, it translocates to the plasma membrane inducing a chloride conductance. Our previous studies comprehensively demonstrated tmCLIC1’s contribution to the progression of GB both *in vitro* and *in vivo*^33,34^. In addition, its specific localization and enrichment to the GSC plasma membrane render tmCLIC1 a valuable pharmacological target for GB treatment.

Here, by combining structural and functional assays *in vitro* and *in vivo* we unveil tmCLIC1 as a privileged metformin membrane interactor in GSCs.

## Results

### Metformin affects GSCs proliferation via tmCLIC1

To assess the interaction between metformin and tmCLIC1, three patient-derived GSC-enriched primary cultures (GBM1-3) and one murine glioma cell line (GL261) lacking CLIC1 expression have been generated through CRISPR-Cas9 technology. Once validated (Extended Data Figure 1), the clones have been rescued with two different CLIC1 forms: (i) the wild type (WT) and (ii) the arginine-to-alanine 29 (R29A) mutant, which is supposed to affect the putative metformin binding site to tmCLIC1, as hypothesized from previous crystallography studies^31^. Extended Data Figure 1 reports the expression level (first two rows from the left) quantification of tmCLIC1 in the plasma membrane (central row) and the ion conductance value (right row), in CRISPR-Cas9 negative control (NC), *Clic1^-/-^*, and rescued for three GBM clones (a, b, and c) plus murine GBM cells GL261(d).

The proliferation rate of these cell populations has been tested in 2D and in 3D models, over 96 hours in the absence or presence of 5 mM metformin (Figure 1 and Extended Data Figure 2). After 96 hours, the drug reduced the proliferation of NC GBM1 cells by approximately 50% (Figure 1a). *Clic1^-/-^* cells showed a slowed-down proliferation rate like metformin-treated NC cells and are insensitive to metformin treatment. Re-expression of WT or R29A CLIC1 protein fully recovered proliferation in the absence of metformin. Strikingly, WT rescued cells were as sensitive to metformin as NC cells (metformin-treated NC vs metformin-treated *Clic1^-/-^* +*CLIC1* WT, n.s. p value= 0.5683) while R29A rescued cells were totally insensitive towards metformin (metformin-treated NC vs metformin-treated *Clic1^-/-^* +*Clic1* R29A rescued, *p value= 0.0164). Similar behavior was observed in 3D cultures (Extended Data Figure 2a) in which metformin incubation reduces the growth of spheroid developed from NC and WT rescued cells. On the contrary, metformin did not exert any effect on KO and the R29A rescued populations. In addition, the antiproliferative activity of metformin was superimposable to that caused by tmCLIC1omab®, a monoclonal antibody targeting the transmembrane form of CLIC1 protein only (Extended Data Figure 2b and c), suggesting that the antiproliferative action of metformin is dependent on tmCLIC1 impairment rather than a direct effect of metformin itself. Worth noting, tmCLIC1omab® was able to decrease proliferation in both WT and R29A mutants, indicating that metformin and tmCLIC1 antibodies don’t share the same epitope target on CLIC1.

**Figure 1.**
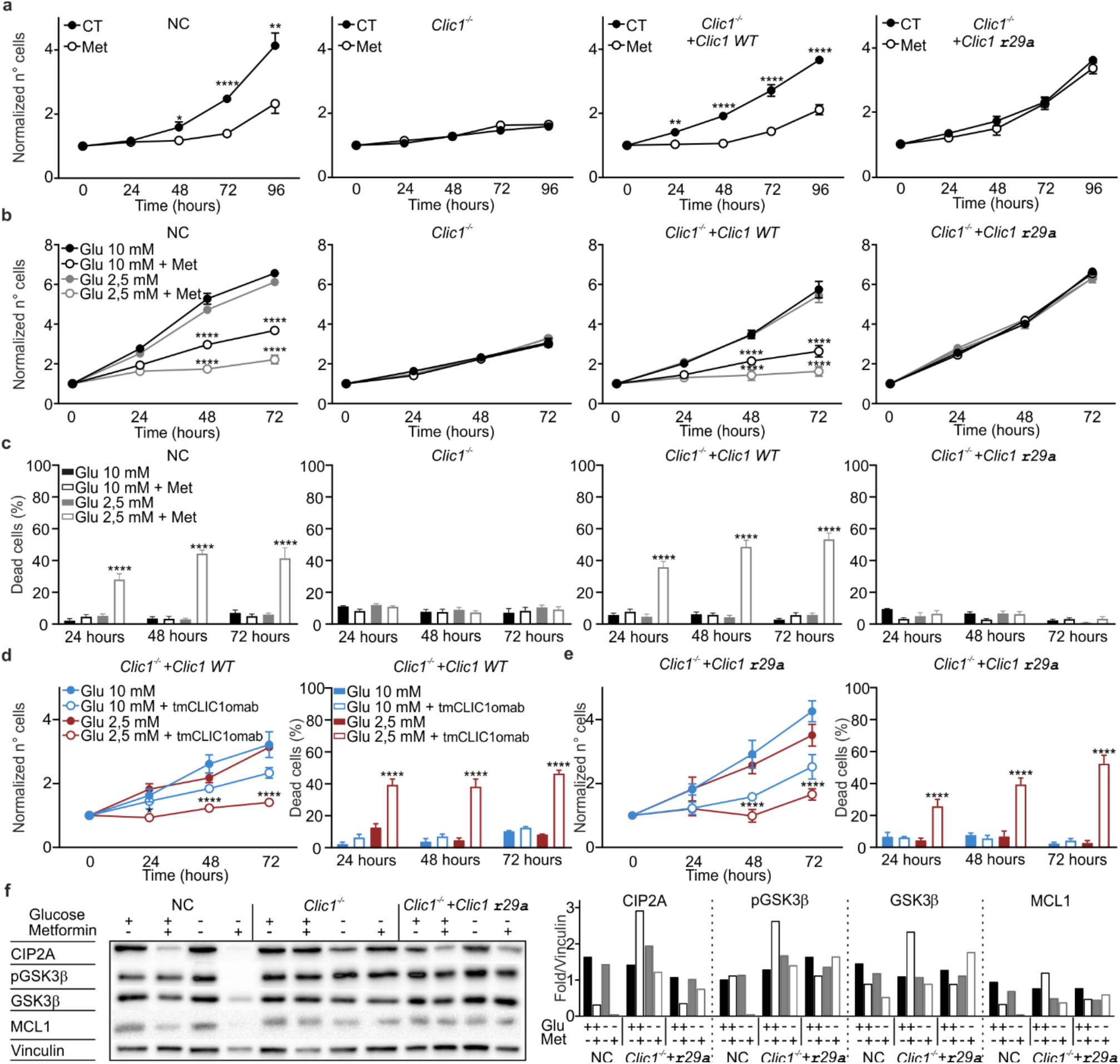
CLIC1 is responsible for the impairment of proliferation caused by metformin. **(a):** Growth curves of GBM1 GSCs over 96 hours in the absence (CT, black circles) or presence (MET, empty circles) of 5 mM metformin treatment in NC, *Clic1^-/-^*, *Clic1^-/-^*+*Clic1* WT, and *Clic1^-/-^* +*Clic1* R29A. NC (CT: 24-96h n=4, Met: 24-96h n=7. 48h *P= 0.0110, 72-96h ****P<0.0001), *Clic1^-/-^* (CT: 24-72h n=4, 96h n=2; Met: 24-72h n=7, 96h n=3), *Clic1^-/-^*+*Clic1* WT (CT: 24, 48, 96h: n=8, 72h n=7; Met 24h n=5, 48-96h n=6. 48h **P=0.0018, 48-96h ****P<0.0001), and *Clic1^-/-^*+*Clic1* R29A (CT: 24, 96h n=8, 48, 72h n=5; Met 24h n=5, 48-96h n=6), mean ± SEM, unpaired t-test. **(b):** Growth curves of GBM1 GSCs over 72 hours in high (black) and low (grey) concentration of glucose and in the absence (filled circles) or presence (empty circles) of 5 mM metformin treatment in NC, *Clic1^-/-^*, *Clic1^-/-^*+Clic1 WT, and *Clic1^-/-^*+Clic1 R29A. NC (n=4; 48-72h ****P<0.0001), *Clic1^-/-^* (n=4), *Clic1^-/-^*+Clic1 WT (n=7, 48-72h ****P<0.0001), Clic1^-/-^+Clic1 R29A (n=6), mean ± SEM, one-way ANOVA, Tukey’s multiple comparison test. **(c):** Analysis of the percentage of dead cells shown in panel b. NC (n=4; 24-72h ****P<0.0001); *Clic1^-/-^*+Clic1 WT (n=7, 24-72h ****P<0.0001), mean ± SEM, one-way ANOVA, Tukey’s multiple comparison test. **(d):** (left) Growth curves of GBM3 GSCs over 72 hours in high (blue) and low (dark red) concentration of glucose and in the absence (filled circles) or presence (empty circles) of 3,5 μg/ml tmCLIC1omab antibody in *Clic1^-/-^* +Clic1 WT (n=3, 24h *P=0.0327 48-72h ****P<0.0001); mean ± SEM, one-way ANOVA, Tukey’s multiple comparison test. (right) Analysis of the percentage of dead cells shown in left panel (n=3; 24-72h ****P<0.0001); mean ± SEM, one-way ANOVA, Tukey’s multiple comparison test. **(e):** (left) Growth curves of GBM3 GSCs over 72 hours in high (blue) and low (dark red) concentration of glucose and in the absence (filled circles) or presence (empty circles) of 3,5 μg/ml tmCLIC1omab antibody in *Clic1^-/-^*+Clic1 R29A (n=3, 48-72h ****P<0.0001); mean ± SEM, one-way ANOVA, Tukey’s multiple comparison test. (right) Analysis of the percentage of dead cells shown in left panel (n=3; 24-72h ****P<0.0001); mean ± SEM, one-way ANOVA, Tukey’s multiple comparison test. **(f):** Representative Western Blot analysis of the expression of CIP2A-GSK3b-MCL1 protein axis in NC, *Clic1^-/-^*, and *Clic1^-/-^*+*Clic1* R29A samples after 72h of culture with or without metformin in high or low concentration of glucose (left). Quantification of CIP2A, phosphorylated form of GSK3b (pGSK3b), GSK3b, and MCL-1 expression in the same cell lines of Western Blot cultured in high (black filled column) and low (grey filled column) concentration of glucose in the absence (black empty column) or presence (grey empty column) of 5mM of metformin.

As reported in our previous research^35^, metformin cooperates with low glucose to induce cell death phenotype in several solid tumors through the activation of the CIP2A-GSK3beta-MCL1 axis, where CIP2A downregulation is directly due to metformin treatment. We, therefore, decided to test if GBM1-3 are sensitive to a combination of metformin and low (2.5 mM) glucose by measuring proliferation rate and cell death. As hypothesized, glioblastoma NC cells clearly showed a switch from a cytostatic to a cytotoxic effect of metformin in a glucose-dependent manner, as described in other solid tumors. Interestingly both *Clic1^-/-^* and R29A rescued cell lines were completely resistant to metformin also in the presence of low glucose cytotoxic combination, elucidating the importance of tmCLIC1 in mediating the cytotoxic effect of metformin (Figure 1b and c). This phenotype was conserved in GBM2, GBM3, and GL261 cells (Extended Data Figure 2g-i). Moreover, tmCLIC1omab® treatment completely phenocopied the effect of metformin when combined to low glucose in both WT and R29A rescued cell lines (Figure 1d and e).

We then explored if the combination of metformin and low glucose in GBM1 cells activated the pathway described in Elgendy, 2019^35^. As expected, metformin with low glucose induced the dephosphorylation of GSK3beta and the subsequent degradation of MCL1 by reducing the key protein CIP2A, in agreement with increased cell death observed upon this treatment (Figure 1f). Strikingly, in *Clic1^-/-^* and R29A rescued cell lines metformin fails to reduce CIP2A, thus downstream phosphorylation of GSK3beta (Ser9) and MCL1 levels resulted unaffected (Figure 1f). On the contrary, R29A rescued cells show an impairment of CIP2A-GSK3beta-MCL1 axis in presence of tmCLIC1omab^®^, demonstrating a key role for tmCLIC1 in this pathway (Extended Data Figure 2l).

Lastly, as a further control, we rescued *Clic1^-/-^* cells with a point mutation in Lys37 (K37A), the only other positive-charged amino acid in tmCLIC1 transmembrane amino acid stretch in addition to Arg29. Despite the mutation, cells have shown sensitivity towards metformin treatment (Extended Data Figure 2d-f), highlighting the importance of Arg29 as the specific residue required for metformin binding and therefore activity. Moreover, GBM1 *Clic1^-/-^* and R29A rescued cells treated with IACS and niclosammide, two inhibitors of mitochondrial complex 1, have demonstrated a strong cytotoxicity (Extended Data Figure 2 j-k).

### Metformin accumulates GSCs in the G1 phase

We previously demonstrated that tmCLIC1 functional activity is subject to the timing of signals that control its membrane insertion and removal^34^. Particularly, its activity is required during the G1/S cell cycle transition. Therefore, we investigated the genetic background impact of the four cell lines on timing of G1/S transition by looking at Cyclin E1 expression levels. This protein accumulates at the G1/S phase boundary and is dramatically degraded as cells progress through the S phase. Cyclin E1 dynamics followed the same trend in all the clones, except for *Clic1^-/-^* cells (Extended Data Figure 3a), displaying a delayed cell cycle progression consistent with a slow-down in cell proliferation, as shown in Figure 1. As expected, metformin-incubated NC and *Clic1* WT rescued cells showed a delayed peak of Cyclin E1. Clic1 R29A rescued cells did not show any timing alteration in cyclin E1 levels after metformin treatment confirming again their insensitivity to the drug (Extended Data Figure 3a). Furthermore, the effect of tmCLIC1omab® was superimposable to that of metformin in NC cells and was conserved in R29A rescued cells (Extended Data Figure 3b). A similar phenotype was observed on GBM2 and GBM3 cells (Extended Data Figure 3c).

### Metformin effect on metabolism depends on CLIC1

As a result of metformin treatment, many cancer cells show strong inhibition of OXPHOS. For this reason, we explored the modulation of GBM cell metabolism (both mitochondrial respiration and glycolysis) upon acute treatment of high concentration (10 mM) metformin in NC, *Clic1^-/-^*, rescued WT, and rescued R29A cells. Both NC and rescued WT cells had similar mitochondrial respiratory rates showing a quick drop in oxygen consumption and a slow rise of extracellular acidification after the injection of metformin, according to a putative inhibition of OXPHOS. On the other hand, *Clic1^-/-^* cells showed a drastic reduction in basal metabolism and seemed totally unaffected by metformin treatment. In addition, rescued R29A cells showed a basal metabolism comparable to rescued WT but were insensitive to metformin treatment (Figure 2a, for GBM1 and Extended Data Figure 4a and c, for GBM2 and GBM3). A comparable effect was observed in cells treated with tmCLIC1omab®, demonstrating that the disruption of tmCLIC1 functional activity quickly impacts mitochondrial respiration and was almost superimposable to metformin treatment (Extended Data Figure 4e-g, GBM2 and GBM3).

**Figure 2.**
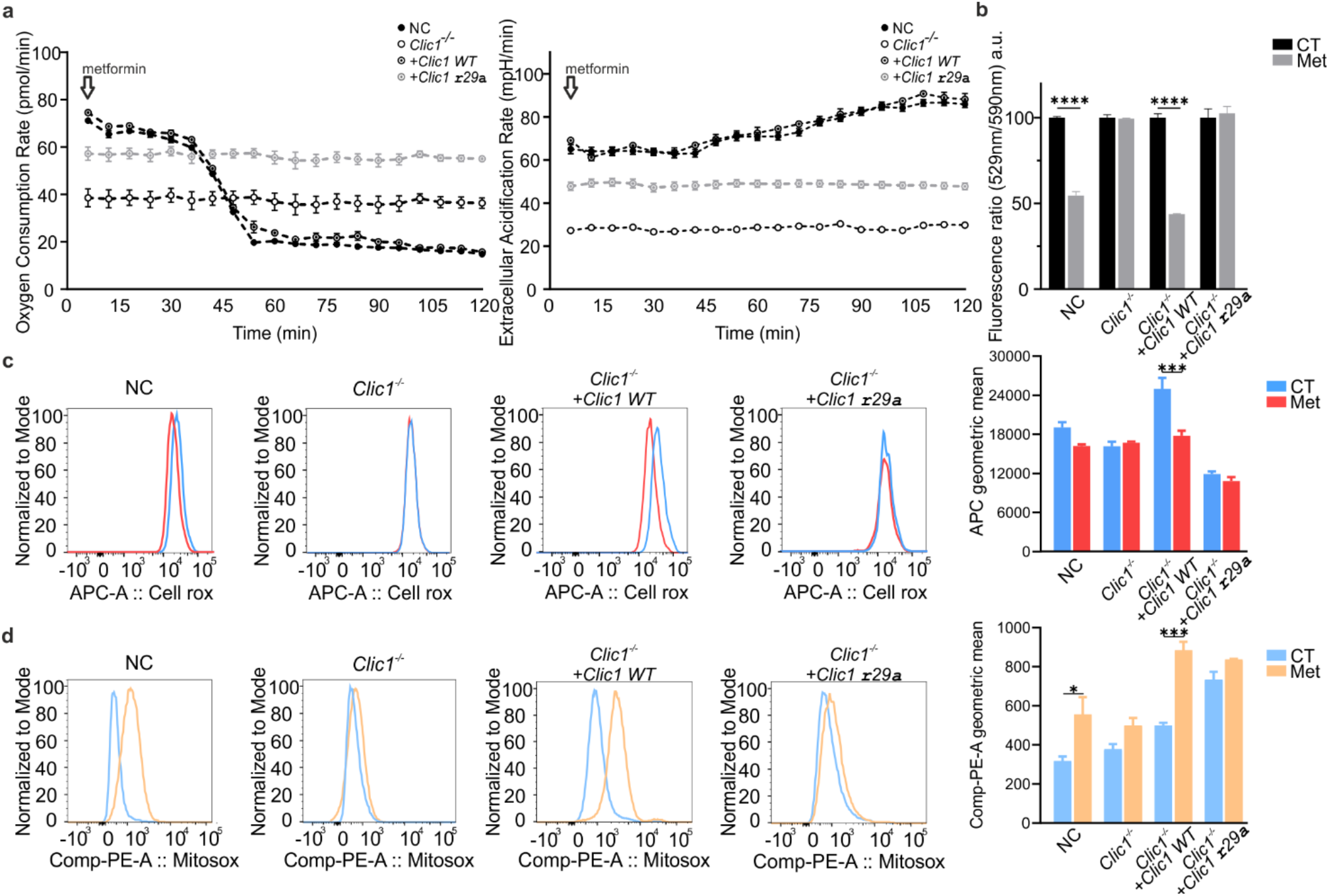
Metformin’s effect on metabolism depends on CLIC1. **(a):** Oxygen consumption rate (left) and Extracellular acidification rate (right) in GBM1 as an indicator of oxidative phosphorylation and glycolitic activity after acute injection of 10 mM metformin in NC (black circles), *Clic1^-/-^* (empty circles), *Clic1^-/-^* +Clic1 WT (black circle dot), *Clic1^-/-^* +Clic1 R29A cells (gray circle dot). Each experimental point was taken every 6 minutes. **(b):** Mitochondrial membrane potential measured in GBM1 through JC-1 probe in NC, *Clic1^-/-^, Clic1^-/-^* +Clic1 WT, and *Clic1^-/-^* +Clic1 R29A in the absence (black columns) or presence (grey columns) of 5 mM metformin treatment. n=2; ****P<0.0001; mean ± SEM, two-way ANOVA, Sidak’s multiple comparison test **(c):** (left) Representative plot of FACS analysis of CellROX^™^ Deep Red Reagent fluorescence for oxidative stress detection in GBM3. Cells were incubated for 3 hours in the absence (blue) or presence (red) of 5mM of Metformin. (right) Mean Fluorescence Intensity of CellROX^™^ Deep Red Reagent. n=3 *Clic1^-/-^* +Clic1 WT: CT vs Met; ***P=0.0002; mean ± SEM, one-way ANOVA, Tukey’s multiple comparison test. **(d):** (left) Representative plot of FACS analysis of MitoSOX^™^ Red Indicator fluorescence for Mitochondrial Superoxide detection in GBM3. Cells were incubated for 3 hours in the absence (blue) or presence (yellow) of 5mM of Metformin. (right) Mean Fluorescence Intensity of MitoSOX^™^ Red Indicator. n=4 NC: CT vs Met; *P=0.0361, *Clic1^-/-^* +Clic1 WT: CT vs Met; ***P=0.0005; mean ± SEM, one-way ANOVA, Tukey’s multiple comparison test.

To further support that the interaction between metformin-tmCLIC1 acts as a trigger of a stress cascade connecting tmCLIC1, mitochondrial activity, and proliferation rate in GBM, we measured mitochondrial membrane potential, using a JC1 probe, 30 minutes after metformin treatment. NC and rescued WT quickly reacted to metformin showing a decrease in mitochondrial potential, while *Clic1^-/-^* and R29A were still unaffected by the treatment (GBM1: Figure 2b and GBM2 and GBM3: Extended Data Figure 4b and d). In addition, to examine whether metformin has an impact on oxidative metabolism, we verified intracellular reactive oxygen species (ROS) and mitochondrial superoxide (O_2-_) production in the four genetic backgrounds of GBM1-3, in the absence or presence of metformin (Figure 2c–d and Extended Data Figure 4h-k). Acute metformin treatment reduced ROS production in NC and WT rescued populations while was ineffective on *Clic1^-/-^* and R29A rescued cells (Figure 2c). Similarly, *Clic1^-/-^* and R29A rescued clones showed no effect of metformin on O_2-_ generation, while in the presence of tmCLIC1 protein we observed a robust increase in superoxide production as a putative signal of mitochondrial acute stress (Figure 2d).

### Evidence of direct binding between metformin and CLIC1 protein

Arginine 29 within the CLIC1 sequence thus appears as the amino acid residue instrumental for the metformin antiproliferative effect. To provide evidence of direct binding between metformin and tmCLIC1, we performed perforated patch clamp experiments, assessing – over time – the whole cell current after perfusion of metformin and tmCLIC1 inhibitor IAA94. Metformin treatment inhibited whole-cell current on NC and WT rescued cells (Extended Data Figure 4a). Conversely, metformin perfusion resulted in a totally ineffective R29A rescued cell current, suggesting metformin’s inability to reach its target. We then opted for a similar approach, but at the single molecular level in single channel outside-out experiments (See Material and Methods section). Strikingly, tmCLIC1 single-channel current, which is blocked by tmCLIC1 inhibitor IAA94, was also blocked after metformin administration in CLIC1 WT rescued cells (Figure 3a; Extended Data Figure 5). On the contrary, the single channel activity of R29A rescued cells was not altered by metformin perfusion, suggesting tmCLIC1 as a direct interactor of metformin in GSCs (Figure 3a; Extended Data Figure 5). To further demonstrate metformin direct binding to tmCLIC1, WaterLOGSY (Water-Ligand Observation with Gradient Spectroscopy)^36,37^ and T_2_ filter (transverse relaxation filter)^38^ NMR signals of metformin (2 mM) were assessed in the absence of cells or in the presence of *Clic1^-/-^*, *Clic1^-/-^* +Clic1WT, or *Clic1^-/-^* +Clic1R29A cells (Figure 3b and c). A remarkable binding effect of metformin in the presence of WT rescued cells was observed, whereas a lower binding effect was observed in the presence of *Clic1^-/-^* cells. Even though the modest metformin binding to *Clic1^-/-^* cells can be ascribed to the existence of other metformin biological targets, the low binding effect recorded also in presence of R29A rescued cells suggests that this specific mutation severely affects metformin binding to CLIC1. These data were further supported by WaterLOGSY and T_2_ filter NMR experiments performed on the double cellular concentration that confirmed a binding effect dependent on cell number (extended Data Figure 5e). NMR experiments were further performed on isolated WT and R29A-mutated recombinant proteins (See Material and Methods section) to confirm the direct molecular interaction of metformin with CLIC1. WaterLOGSY and T_2_ filter NMR experiments showed that metformin binds to the WT form of CLIC1 and not to the R29A mutated form (T=25 °C, extended Data Figure 5d, e). These data confirm that the mutation of a charged hydrophilic residue (R) in position 29 into a hydrophobic one (A) strongly affects metformin binding to CLIC1 at molecular resolution.

**Figure 3.**
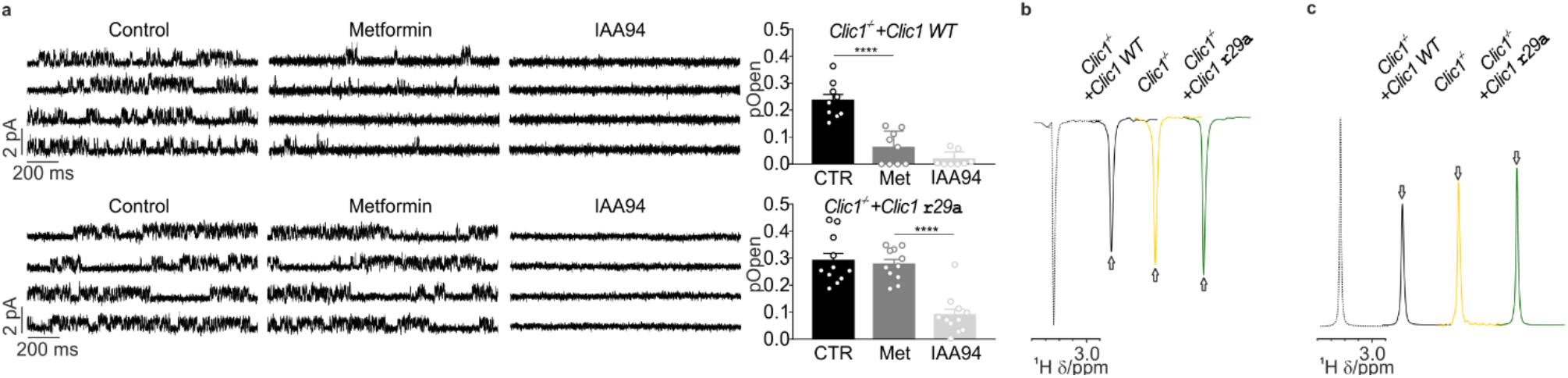
Metformin directly interacts with tmCLIC1. **(a):** (Upper panel) Representative traces of GBM3 *Clic1^-/-^*+*Clic1* WT GSCs outside-out experiments during the perfusion of the vehicle (CTR) or the compounds, as indicated (left). Quantification of tmCLIC1 single channel open probability in the mentioned conditions (right). CTR and Met n=9, IAA94 n=8; ****P<0.0001; mean ± SEM, one-way ANOVA, Tukey’s multiple comparison test. (Bottom panel) Representative traces of GBM3 *Clic1^-/-^+Clic1* R29A GSCs outside-out experiments during the perfusion of the compounds as indicated (left). Quantification of tmCLIC1 single channel open probability in the mentioned conditions (right). n=11; ****P<0.0001; mean ± SEM, one-way ANOVA, Tukey’s multiple comparison test. **(b):** WaterLOGSY and **(c):** T_2_ filter 1^H^ NMR spectra of 2 mM metformin in absence (dashed line) or presence of *Clic1^-/-^* (yellow), *Clic1^-/-^*+Clic1 WT (black), and *Clic1^-/-^*+Clic1 R29A (green), cells. Arrows highlight signal changes showing a significant effect of metformin binding to cells in the presence of *Clic1 ^-/-^*+Clic1 *WT* compared to *Clic1^-/-^*+Clic1 R29A and *Clic1^-/-^*.

### Metformin affects tumor expansion in zebrafish embryos and mice glioblastoma models

To assess whether the cytostatic/cytotoxic effect of metformin is maintained in more complex and *in vivo* systems, zebrafish embryos were orthotopically injected with GBM1 cells at 48 hours post fertilization and the tumor mass was measured after 72 hours in the absence or presence of metformin diluted in embryos’ water (5 mM). The results depicted in Figure 4a, and b are consistent with our *in vitro* observations (Figure 1), metformin effectively reduced tumor expansion in xenografts obtained from NC and *Clic1^-/-^* WT rescue cells (Figure 4a, b), while its ability was totally lost when tmCLIC1 function in GSCs was impaired (*Clic1^-/-^*) or in the presence of R29A point mutation (Figure 4a and b).

**Figure 4.**
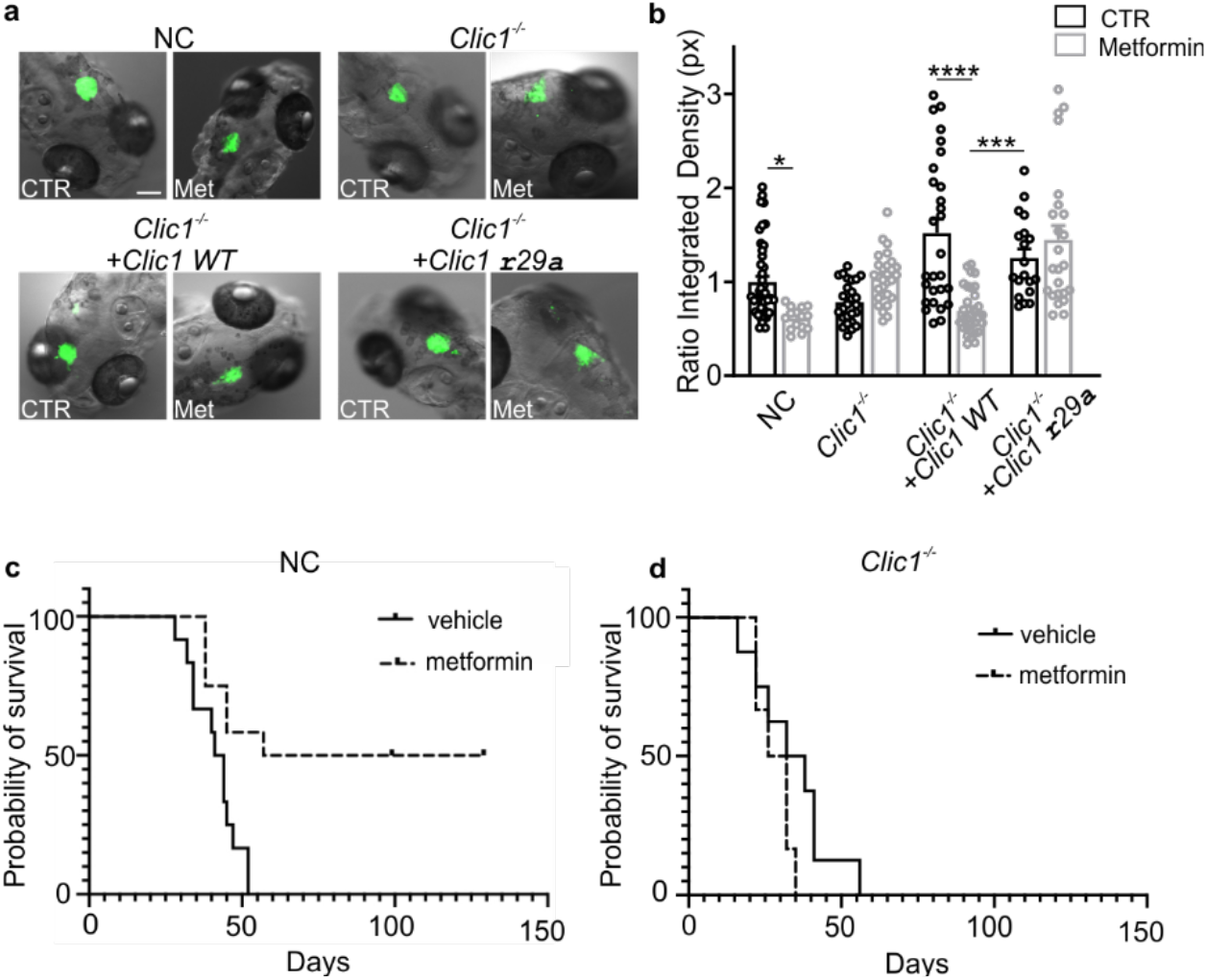
Metformin impairs the growth of patient-derived xenografts in zebrafish embryo models and mouse glioblastoma in immunocompetent mice. **(a):**. Representative pictures showing the expansion of the tumor mass at 72 hours post-injection in zebrafish embryos’ brain injected with NC, *Clic1^-/-^, Clic1^-/-^* +Clic1 WT, *Clic1^-/-^* +Clic1 R29A cells GBM1 cells in absence or presence of metformin 5 mM (as indicated) dissolved in embryos’ water. Scale bar 100 μM. **(b):** Quantification of the integrated density of the tumor mass in absence (black) or presence (gray) of metformin. Every experimental point represents the expansion of the tumor mass measured in the single embryo’s brain. NC: CT n=44, Met n=17; *P<0.0262; *Clic1^-/-^*+Clic1 WT: CT n=28, Met n=31; ****P<0.0001; *Clic1^-/-^*+Clic1 R29A; CT n=19; **P=0.0025; mean ± SEM, one-way ANOVA, Tukey’s multiple comparison test. **(c):** Survival curve (vehicle, n=12 and Metformin n=12; P = 0.0035, log-rank test) of metformin or vehicle-treated GL261 WT mice. **(d):** Survival curve (vehicle, n=8 and metformin n=6; P = 0.1403, log-rank test) of metformin or vehicle-treated GL261 Clic1^-/-^.

To confirm these results in an *in vivo* mammalian setting, we then took advantage of GL261 cells to generate a syngeneic murine GBM model ^39^. The administration of metformin (10 mM) in drinking water for up to 100 days, allowed measurable brain concentrations (reaching a steady state of about 12 +/- 0.4 nM, n=8) and effectively prolonged the survival of mice orthotopically injected with GL261 WT cells (p=0.0035) and reduced tumor incidence (vehicle: 12/12 mice; metformin 6/12 mice) (Figure 4c). Conversely, metformin was ineffective in extending survival in mice bearing GL261 *Clic1^-/-^-derived* tumors (p=0.1403) (Figure 4d).

## Discussion

Glioblastoma is still one of the deadliest solid tumors with a high recurrence rate after standard of care. In the perspective to set a more effective therapy, it is imperative to hit the pool of GSCs. Targeting preferentially the subpopulation of GSCs would be instrumental to eradicate tumor driving force, affecting GBM resistance to conventional therapy, and hamper tumor relapse. GSCs show significantly higher levels of tmCLIC1 compared to tumor bulk cells^31^ and normal brain tissue^33^. Thus, drugs aiming at tmCLIC1 blockade would discriminate and target preferentially GSCs and produce fewer side effects on the central nervous system.

The present investigation demonstrates that tmCLIC1 is the main membrane interactor for metformin in GBMM cells. Failure of metformin binding to tmCLIC1 prevents the antineoplastic effect of the biguanide compound *in vitro* as well as *in vivo*. Our results are in accordance with previous studies describing the effect of metformin on downstream metabolic pathways. Thus, metformin reacts with tmCLIC1, causing a fast depolarization of cellular membranes with a quick reaction of mitochondria at the point of oxygen consumption and membrane potential. We hypothesize that this metformin-mediated stress starts from tmCLIC1 inactivation by direct binding, goes through a fast metabolic adaptation and mitochondrial dysfunction, and leads to ROS production, ultimately affecting protein synthesis and proliferation, which results in cell cycle arrest^40^. The ability of *Clic1^-/-^* and R29A mutant to revert both cytostatic and cytotoxic effects of metformin on GBM, strongly supports the idea of tmCLIC1 as a *bona fide* metformin primary interactor. Furthermore, we elucidated the requirement of a single amino acid residue (R29A) for metformin binding to CLIC1 protein. We observed a strong dependency between metformin and CLIC1 protein when interaction is disrupted (in by expression of the mutant version or the absence of CLIC1) cells not only become resistant to metformin but no metabolic perturbations are observable. Additionally, when combined with low-glucose, metformin’s effect switches from cytostatic to cytotoxic because of the impairment of the CIP2A-GSK3beta-MCL1 axis. Understanding the key mechanisms behind tmCLIC1 control of these pathways would represent a crucial step in the knowledge of GBM progression, unveiling novel possible therapeutic targets.

Our investigation stated that *in vitro*, the metformin antiproliferative effect is exerted at a millimolar concentration range. However, *in vivo* experiments demonstrate that mice in which metformin was administered through drinking water exhibit an increase in survival, despite metformin’s brain quantification have been reported to be in the nanomolar range. It is possible that the constant fresh drug circulation, continuously reaching the brain during the prolonged administration in the in vivo treatments (up to 100 days in vivo) allowed a drug steady state compatible with a persistent saturation and inhibition of all the active tmCLIC1 expressed by GBM cells (between 10 and 50 active channels at 0 mV membrane potential). The requirement of lower concentrations *in vivo* than *in vitro*, is also an important information to translate these studies in clinics, in which the already available extended release formulation may provide the same sustained activity on GBM cells than the administration of the drug in the mice drinking water.

In a recent publication, we demonstrated the existence of GBM subsets whose aggressiveness is unrelated to CLIC1 expression whereas displaying resistance towards metformin treatment^42^. This introduces the possibility to establish novel eligibility criteria for discriminating patients’ cohorts in a personalized trial setting based on CLIC1 expression.

Based on these results, future investigations aiming at enhancing metformin-tmCLIC1 interaction will improve the ability to target unambiguously glioblastoma stem cells.

## Acknowledgment

This work has been supporter by AIRC IG Grant to MM. We are thankful to Stefania Castiglione and Yuli Buckley for technical assistance.

## AUTHOR CONTRIBUTIONS

IV, FC, and RC conducted the majority of experiments including, generic engineering, western blots, fluorescence assays, electrophysiological experiments, metabolic assays, cytofluorimetric analysis, growth curves experiments, zebrafish embryos trials, and data analyses. GC contributed to the recombinant protein isolation. MR and LP helped with electrophysiology recordings and zebrafish trials. CT assists to cytofluorimetric assays. MV, BB, and SG performed NMR binding experiments. CR, SF, DO, and GP execute all the experiments in mice model. GA and GP operate metformin measurements in mice brains. IV, FC, RC, TF, SM, and MM prepared the manuscript. GC, CR, SF, DO, CT, SG, FB, AF, AC, GP participated in data analysis and manuscript writing. TF, SM, and MM direct manuscript and figures editing. IV, FC, TF, SM, and MM conceived and supervised the project.

## Methods

### Reagents

Indanyloxyacetic acid 94 (IAA94) (Sigma-Aldrich) was used to specifically inhibit CLIC1 activity. Initial concentration: 50 mM dissolved in absolute ethanol. Final concentration: 100 μM.

1,1-Dimethylbiguanide hydrochloride (metformin) (Sigma-Aldrich) is used as an alternative CLIC1 inhibitor. Initial concentration: 1M dissolved in ultrapure deionized water. Final concentration 5 mM or 10mM.

PD033 isethionate (Sigma-Aldrich) is an inhibitor of cyclin-dependent kinase (CDK) 4 and 6. It was used to synchronize cells in G1 phase of the cell cycle at a concentration of 2.5 μg/ml.

Niclosamide (Sigma-Aldrich) was reported to be an uncoupler of oxidative phosphorylation. Initial concentration: 10mM dissolved in DMSO. Final concentration 4 μM.

IACS-010759 (Sigma-Aldrich) was used as mitochondria complex I blocker. 10mM dissolved in DMSO. Final concentration 100 nM.

### Cell cultures

Human glioblastoma stem cells (GSCs) primary cultures, named as GBM1, GBM2, and GBM3 were isolated and maintained as previously described^43^. GBM1 GSCs grow in suspension as spheroid aggregates. Twice a week neurospheres were mechanically dissociated into single-cell suspension to form secondary neurospheres. GBM2 and GBM3 GSCs grow in adhesion on plastic supports. For some experiments, GSCs were grown also on plates coated Geltrex LDEV-Free hESC-qualified Reduced Growth Factor Basement Membrane Matrix (Thermo Fisher Scientific).

### *Clic1^-/-^* mutant generation by CRISPR-Cas9 technology

Patient-derived GSCs were transfected with transEDIT lentiviral gRNA plus Cas9 expression (pCLIP-All-hCMV-ZsGreen V66) lentiviral vector according to manufacturer’s protocol (Transomic). Two plasmids were used, a gRNA targeting a specific region of *Clic1^-/-^* coding sequence (TGAGTGCCCCTATACCTGGG) and one targeting GFP gene as negative control (NC). Plasmids carry ZsGreen fluorophore as a selection marker.

### Plasmids

For *in vitro* experiments CLIC1wt-pIRES2-EGFP or CLIC1R29A-pIRES2-EGFP plasmids have been used to rescue transiently *clic1^-/-^* cells. For *in vivo* experiments, *Clic1* in its WT or R29A mutated form have been cloned into pCDH cDNA and used to rescue stably *clic1^-/-^* cells using a lentiviral transduction protocol.

### OCR and ECAR measurements

Measurement of OCR and ECAR have been performed using the seahorse XF96 (Agilent), experimental conditions have been plated in 5 replicates and experiments followed manufacturer’s protocol. To test cells metabolic adaptation upon metformin acute administration, 10mM metformin have been injected by the instrument, OCR and ECAR have been measured up to 120 minutes.

### Mitochondrial potential analysis

Cells have been plated 2×10^4^ GBM1 cells or 7×10^3^ GBM2-GBM3, 5mM metformin and JC1 probe (Thermofisher) have been added together to the cells and incubated 30 minutes. Fluorescence have been measured by BD FACSCelesta^™^ as described in manufacturer’s protocol.

### Quantification of intracellular ROS and mitochondrial SOX

GBM1-3 and GL261 cells have been incubated for 3 hours in absence or presence of 5 mM metformin and stained with CellROX^™^ deep red reagent and MitoSOX^™^ red mitochondrial superoxide indicator according to manufacturer’s instructions. Cells were analyzed by BD FACSCanto^™^ II. Data have been analyzed using FlowJo Single Cell Analysis Software v10, calculating the geometric mean of the histogram resulting from cellular fluorescence.

### Protein extraction and Western Blot analysis

Cells were lysed through the addition of hot Lysis Buffer (0.25M Tris-HCl pH 6.8, 4% SDS, 20% Glycerol in water). Samples were sonicated for 30 minutes, syringed, and boiled for 10 minutes at 95°C. Samples were centrifuged at 4,000 x g for 15 minutes, supernatants were collected and stored at −20°C. Protein concentration was evaluated through BCA assay (Thermo Fisher Scientific). Samples were loaded onto 12% SDS-polyacrylamide electrophoresis gel (PAGE) and run at 100 V in a running buffer. Separated proteins were transferred to a nitrocellulose membrane (Amersham Protran, GE Healthcare) with of 0.45 μm pore size at 100 V for 1 h on ice in transfer buffer. After blocking, membranes were incubated with primary antibodies diluted in PBS/BSA 5% overnight at 4°C, then washed three times to remove the excess of primary antibodies and incubated 1 h RT with secondary antibody solutions. Finally, membranes were incubated with SuperSignal® West Femto Maxium Sensitivity Substrate (Thermo Fisher Scientific) 1 minute in the dark. Immunoreactive protein bands were detected using ChemiDoc Touch® imaging system (BioRad). Results were analyzed using ImageLab software (BioRad).

### Growth Curves

2×10^4^ GBM1 cells or 7×10^3^ GBM2-GBM3 cells were plated in 500 μL of growth medium per well (supplemented with treatment when needed) in a 24-well plate. For each experimental condition and time point, cells were plated in triplicates. Cells were plated in 24-multiwell and counted at given time points to build up the growth curve. Cells were collected and centrifuged, and the resuspended pellet was diluted 1:1 with Trypan Blue. Countess II FL automated cell counter (Thermo Fisher Scientific) was used to count them. Data normalized to their controls.

### 3D Cultures

GBM1 cells (NC, *Clic1^-/-^*, *Clic1^-/-^* +Clic1WT, and *Clic1^-/-^* +Clic1R29A) were plated in 24-well plates at a density of 2×10^4^cells. After the formation of a solid 3D structure (24 to 48 hours after plating) single spheroids were transferred to a new 24-well plate in fresh medium with or without metformin (5mM) and the first photo was captured to measure the initial area. Spheroids were incubated for 72 hours, and pictures were captured to normalize the final area on the respective initial area. Area measured with ImageJ software (Freehand selection).

### Fluorescence intensity assay

Cells (1×10^6^ cells/well) were washed three times and incubated in a blocking solution for 30 minutes in ice. Primary antibody solutions were added to samples and incubated for an additional 2h in ice. After washes, samples were incubated with secondary antibody solution for 1h in ice in the dark. Samples were washed three times and distributed into a black 96-well plate. All the washes and staining steps were performed maintaining cells in suspension. Samples were analyzed at Ensight Multimode Plate Reader (PerkinElmer’s) using the appropriate filter to visualize fluorescence intensity emitted by Alexa Fluor 555 conjugated antibody. Fluorescence intensity values of samples incubated with tmCLIC1omab® (3.5 μg/ml) are proportional to the amount of tmCLIC1 and were normalized to values of samples incubated only with secondary antibody (1:400, Donkey anti-mouse conjugated to Alexa Fluor 555, Thermo Fisher Scientific).

### Patch clamp experiments

The voltage step protocol used to isolate current/voltage relationships consisted of 800 ms pulses from −60 mV to +60 mV (20 mV voltage steps). The holding potential was set according to the resting potential of the single cell (between −40 and −80 mV). CLIC1-mediated chloride currents were isolated from other ionic currents by perfusing IAA94 100 μM dissolved in the bath solution and by mathematical subtraction of the residual current from the control.

Outside-out experiments were performed to isolate the single tmCLIC1 channel, maintaining physiological conditions with the opportunity to expose it to metformin perfusion. The whole experimental procedure was composed of two independent protocols. First, the channel was identified by a voltage step protocol from −40 mV to + 40 mV (20 mV voltage steps). Next, within the same experiment, the membrane was clamped at 0 mV and blockers (metformin followed by IAA94) were consequently perfused after at least 3 minutes of recording.

In time-course experiments, the holding potential was set according to the resting potential of the single cell and every 5 seconds a +60 mV voltage step was applied. The current was measured at the end of the 800 ms voltage step. Once the current amplitude reached a constant value metformin (5mM) and/or (IAA94) were perfused.

Analysis was performed using Clampfit 10.2 (Molecular Devices) and OriginPro 9.1.

The solutions used were the following:

Bath solution (whole cell and outside-out experiments): NaCl 140 mM, KCl 5 mM, Hepes 10 mM, glucose 5 mM, CaCl_2_ 2 mM, MgCl_2_ 1 mM, pH 7.4.
Pipette solution (whole cell): KCl 135 mM, NaCl 5 mM, Hepes 10 mM, MgCl_2_ 1 mM, CaCl_2_ 2 mM, Gramicidin 2.5 μg/ml, pH 7.4.
Pipette solution (outside-out): KGluconate 120 mM, KCl 20 mM, TEACl 5mM, Hepes 10 mM, CaCl_2_ 0.1 mM, pH 7.

### Cyclin E1 expression

Cells were synchronized in early G1 by incubating them with PD0332991 – a CDK4/6 inhibitor effective at arresting cells in the early G1 phase – for 24 hours and released for the appropriate time in the absence or presence of 5 mM metformin.

### Recombinant protein purification

To express recombinant CLIC1 protein, Rosetta (DE3) *E. coli* cells were transformed with pET_2_8a-CLIC1 plasmid, let grow in continuous shaking (270rpm), and induced with 1mM IPTG once the bacterial culture reached an OD = 1. Protein expression was carried on for 3 hours from the induction, at the same shaking speed. After that interval, cells were collected and lysed by sonication and the soluble fraction was isolated through centrifugation (30 minutes, 4°C, 16000 x g).

The soluble fraction was passed through a HisTrap Nickel column to isolate our protein of interest carrying His-tag sequence. The obtained eluate was used to perform a second chromatographic run in Superdex 75 column, to separate the recombinant protein from eventual contaminants and to put the protein in its final protein buffer (150mM NaCl, 10mM HEPES, pH 7.4). The resulting concentration of the protein solution was between 1-5 mg/mL. The same procedure was done for both CLIC1-WT and CLIC1-R29A.

### NMR binding experiments

NMR samples were analyzed by a Bruker FT NMR Avance III 600 MHz spectrometer with an automatic sample changer SampleJetTM with temperature control, in phosphate-buffered saline (PBS), pH 7.4, 10% D2O for a lock signal

The metformin stock (Sigma-Aldrich) was freshly prepared before each session of NMR experiments at a concentration of 200 mM in MilliQ water.

#### NMR binding experiments on recombinant purified proteins

50 μM of metformin was tested in the absence and in presence of 10 μM CLIC1wt and 10 μM CLIC1[R29A] recombinant proteins, at 25 °C using a 5 mm SEF (Selective 19F, 1H Decoupling) probe with z-gradient coil. The water suppression in all 1H experiments was achieved with the excitation sculpting sequence^44^. The two-water selective 180° square pulses and the four PFGs of the scheme were 2.5 and 0.8 ms in duration, respectively, and a gradient recovery time of 0.25 ms. The 1D ^1^H and the transverse relaxation filter (T_2_ filter)^45^ were recorded using a spectral width of 20 ppm, the acquisition time of 1.3 s, 7 s of relaxation delay, 32 scans. Two T_2_ filter experiments were recorded for each sample with a CPMG spin-echo train sequence with a total τ of 0.72 s and 1.44 s, respectively. The WaterLOGSY experiments were achieved with a 15 ms long 180° Gaussian-shaped pulse, aq 0.42 s, mixing time of 1.2 s, relaxation delay of 2 s, 1024 scans.

#### In cell NMR binding experiments

Cells (*Clic1^-/-^*, *Clic1^-/-^* +Clic1WT, and *Clic1^-/-^* +Clic1R29A) were grown in T75 flasks until 90-100% confluence was reached (~15×10^6^ cells). After 24 h, cells were washed twice with PBS, detached with Triple (500 μL for each flask) for 2 minutes at 37 °C, resuspended in PBS (2 mL for each flask) and spun at 180 g for 6 minutes. The supernatant was then discarded, and the pellet was resuspended at a final concentration of 7.23×10^6^ cells/ml in deuterated PBS. 450 μL of cells solution was transferred into a 5 mm NMR tube together with 50 μL of a 20 mM metformin stock solution. Final concentrations in the NMR tube were 6.5×10^6^ cells/ml, 2 mM metformin, 500 μM TSP (chemical shift reference) and 10% D2O (lock signal). An NMR tube containing 450 μL PBS and 50 μL of the same 20 mM metformin stock was also prepared as a reference.

All in cell NMR experiments were recorded at 37 °C using a 5 mm CryoProbe QCI 1H/19F-13C/15N-D quadruple resonance, a shielded z-gradient coil, and an automatic sample changer SampleJet NMR system with temperature control. T_2_ filter and WaterLOGSY experiments were recorded with the same parameters optimized for the recombinant proteins; only the spectral width was reduced to 14 ppm.

### Patient-derived orthotopic xenograft in Zebrafish embryos

Wild type zebrafish embryos AB at 24 hours post fertilization (hpf) were soaked in embryo medium with 0.2 mM 1-phenyl 2-thiourea (PTU) and incubated for further 24 h at 28.5 °C. At 48 hpf, the embryos were dechorionated and anesthetized with 0.0003% tricaine prior to injection. Anesthetized embryos were positioned on a wet agarose 1% pad. The hindbrain of each embryo was injected with approximately 150-200 cells (ZsGreen-positive for NC and KO cells and GFP-positive for rescued cells) using an Eppendorf FemtoJet® microinjector combined with a stereomicroscope (MZ APO, Leica). After transplantation, embryos were incubated for 4h at 32° C and checked for the presence of fluorescent cells in the correct site. Then embryos were incubated at 32° C in embryo’s water for the following three days.

For metformin treatment, screened embryos were transferred in a 48-well plate with 5 mM metformin prior to the incubation at 32° C.

On the same day of the injection and at 5 days post-fertilization (3 days after injection) images of the tumors were captured using a fluorescent stereo microscope and the relative integrated density – obtained by the product of the mean pixel fluorescence intensity and the pixel area of the tumoral mass – was calculated as the ratio between the final and the initial tumor integrated density using ImageJ software.

### GBM orthotopic transplantation in mice

The project has been approved by the Italian Ministry of Health (Authorization 845/2019-PR). All animal procedures were approved by the Organismo per il Benessere e Protezione Animale (OPBA) of the Cogentech animal facility and the experiments were performed in accordance with the Italian laws (D.L.vo 116/92 and following additions), enforcing EU 86/609 Directive.

4-6 weeks old female C57BL/6/J mice obtained from Charles River Laboratories were housed at the Cogentech animal facility according to the guidelines set out in Commission Recommendation 2007/526/EC, 18 June 2007. Mice were intraperitoneally anesthetized with tribromoethanol (0.1 ml/10 g of body weight) and GL261 WT and Clic1^-/-^ were resuspended in sterile PBS (10^5^/2uL) to be stereotaxically injected into the mice nucleus caudatus (coordinates from bregma: 1 mm posterior, 3 mm left lateral, and 3.5 mm in depth). Tumor-bearing mice were monitored daily and sacrificed at the appearance of neurological symptoms and/or signs of morbidity or 20% body weight loss. Metformin was administered in the drinking water (10mM), starting from 1 week before cell injection until the experiment endpoint.

### Brain metformin concentration measurement

Gas chromatography-mass spectrometry (GC-MS) analyses were performed using a Hewlett Packard 5890 Series II gas chromatograph equipped with a Hewlett Packard 5971A mass selective detector and cool on-column injector. A 25m×0.2mm×0.3μm cross-linked methyl silicone gum column was used equipped with a deactivated retention gap. The analysis was performed by thermal gradient, increasing the initial oven temperature of 120 °C to 220 °C. Deuterated metformin was used as internal standard, and the ions m/z = 303 (derivatized metformin) and m/z = 309 (derivatized 2H6-metformin) were monitored.

After preparing a calibration curve obtained by derivatizing metformin standard solutions with N-methyl-bis(trifluoroacetamide) (MBTFA), frozen brain samples were homogenized and lyophilized. Accurately weighed amount of lyophilized mouse brain tissue was extracted with methanol, and the methanolic suspension was vortexed and sonicated. The methanolic surnatant underwent solid phase extraction purification, followed by derivatization with MBTFA.

### Statistical analysis

All data were plotted using GraphPad Prism 7 software (GraphPad Software Inc., San Diego, California), by which all mean values and standard errors have been calculated. Statistical analysis of these data has been performed on the same software. To compare data between two different conditions, an unpaired t-test analysis has been used. A one-way ANOVA test was used to compare more than two groups within the same experimental condition. Each condition in all experiments was supported by at least 3 independent replicates. p < 0.05 was considered statistically significant

