## Supplemental File for "Metformin antiproliferative activity is exclusively mediated by the membrane functional expression of the Chloride Intracellular Channel 1 in glioblastoma stem cells"

**Extended Data**

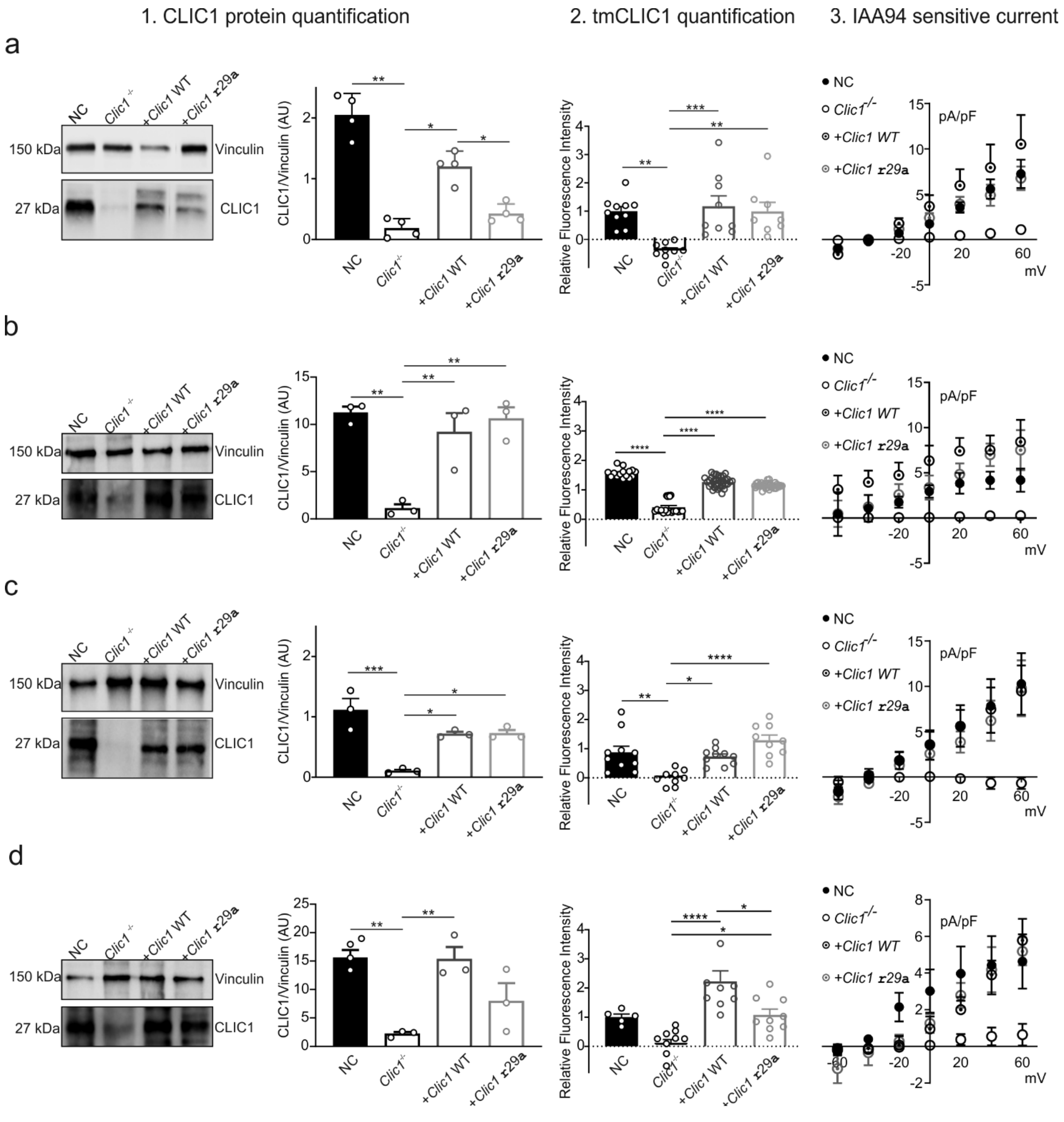


**Extended Data Figure 1 | CLIC1 expression, localization, and quantification in the membrane of GBM1 (a), GBM2 (b), GBM3 (c) primary cultures, and GL261 cells (d).**

First two left columns: representative western blot analyses and relative quantification of CLIC1 protein in NC, *Clic1*$^{-/-}$, and rescued GSCs populations' lysates; mean ± SEM, one-way ANOVA, Tukey's multiple comparison test. **(a):** n=4; NC vs Clic1$^{-/-}$ **P=0.0013; *Clic1*$^{-/-}$+*Clic1* WT vs Clic1$^{-/-}$ *P=0.0366; *Clic1*$^{-/-}$+*Clic1* R29A vs *Clic1*$^{-/-}$+*Clic1* WT *P=0.0355. **(b):** n=3; NC vs Clic1$^{-/-}$ **P=0.0018; Clic1$^{-/-}$+Clic1 WT vs Clic1$^{-/-}$ **P=0.0074; Clic1$^{-/-}$+Clic1R29A vs Clic1$^{-/-}$ **P=0.0028. **(c):** n=3; NC vs Clic1$^{-/-}$ ***P=0.0004; Clic1$^{-/-}$+Clic1 WT vs Clic1$^{-/-}$ *P=0.0108; Clic1$^{-/-}$+Clic1 R29A vs Clic1$^{-/-}$ +Clic1 WT *P=0.0101. d) n=3; NC vs Clic1$^{-/-}$ **P=0.0068; Clic1$^{-/-}$+Clic1 WT vs Clic1$^{-/-}$ **P=0.0077.

Third column from the left: relative fluorescence intensity of tmCLIC1 for a, b, c, d clones; mean ± SEM, one-way ANOVA, Tukey's multiple comparison test.

**(a):** (NC n=10; *Clic1*$^{-/-}$ n=9; *Clic1*$^{-/-}$+*Clic1* WT n=9; *Clic1*$^{-/-}$+*Clic1* R29A n=8). NC vs *Clic1*$^{-/-}$ **P=0.0020; *Clic1*$^{-/-}$+*Clic1* WT vs *Clic1*$^{-/-}$***P=0.0007; Clic1-/-+Clic1 R29A vs *Clic1*$^{-/-}$ **P=0.0040. **(b):** (NC n=15; *Clic1*$^{-/-}$n=19; *Clic1*$^{-/-}$+*Clic1* WT and *Clic1*$^{-/-}$+*Clic1* R29A n=35). ****P<0.0001.

**(c)**: (NC, *Clic1*$^{-/-}$, and *Clic1*$^{-/-}$+*Clic1* WT n=10, *Clic1*$^{-/-}$+*Clic1* R29A n=9). NC vs *Clic1*$^{-/-}$ **P=0.0020; *Clic1*$^{-/-}$+*Clic1* WT vs *Clic1*$^{-/-}$*P=0.0118; *Clic1*$^{-/-}$+*Clic1* R29A vs *Clic1*$^{-/-}$ ****P<0.0001.

**(d)**: (NC n=5; *Clic1*$^{-/-}$ n=10; *Clic1*$^{-/-}$+*Clic1* WT and *Clic1*$^{-/-}$+*Clic1* R29A n=9). *Clic1*$^{-/-}$+*Clic1* WT vs *Clic1*$^{-/-}$****P<0.0001; *Clic1*$^{-/-}$+*Clic1* R29A vs *Clic1*$^{-/-}$*P=0.0228; *Clic1*$^{-/-}$+*Clic1* WT vs *Clic1*$^{-/-}$+*Clic1* R29A *P=0.0116.

Fourth column. Electrophysiology recording in perforated patch clamp of the whole cell current of single glioblastoma stem cells in the four genetic backgrounds; mean ± SEM, one-way ANOVA, Tukey's multiple comparison test.

**(a):** (NC n=7; *Clic1*$^{-/-}$ n=7; *Clic1*$^{-/-}$+*Clic1* WT n=5; *Clic1*$^{-/-}$+*Clic1* R29A n=4); *Clic1*$^{-/-}$+*Clic1* WT vs *Clic1*$^{-/-}$: 0 mV **P=0.0047; 20 mV **=0.0017; 40 mV **=P0.0052; 60 mV **P=0.0056).

**(b):** (NC n=6; *Clic1*$^{-/-}$ n=5; *Clic1*$^{-/-}$+*Clic1* WT n=6; *Clic1*$^{-/-}$+*Clic1* R29A n=4); *Clic1*$^{-/-}$+*Clic1* WT vs *Clic1*$^{-/-}$: 0 mV ****P<0.0001; 20 mV **=0.0017; 40 mV **P=0.0019; 60 mV *P=0.0204; *Clic1*$^{-/-}$+*Clic1* R29A vs *Clic1*$^{-/-}$: 0 mV **P=0.0050; 40 mV **P=0.0086).

**(c)**: (NC n=8; *Clic1*$^{-/-}$ n=5; *Clic1*$^{-/-}$+*Clic1* WT n=6; *Clic1*$^{-/-}$+*Clic1* R29A n=5).

**(d)**: (NC n=5; *Clic1*$^{-/-}$n=5; *Clic1*$^{-/-}$+*Clic1* WT n=4; *Clic1*$^{-/-}$+*Clic1* R29A n=5); *Clic1*$^{-/-}$+*Clic1* WT vs *Clic1*$^{-/-}$:60 mV *P=0.0205; *Clic1*$^{-/-}$+*Clic1* R29A vs *Clic1*$^{-/-}$: 60 mV *P=0.0312.

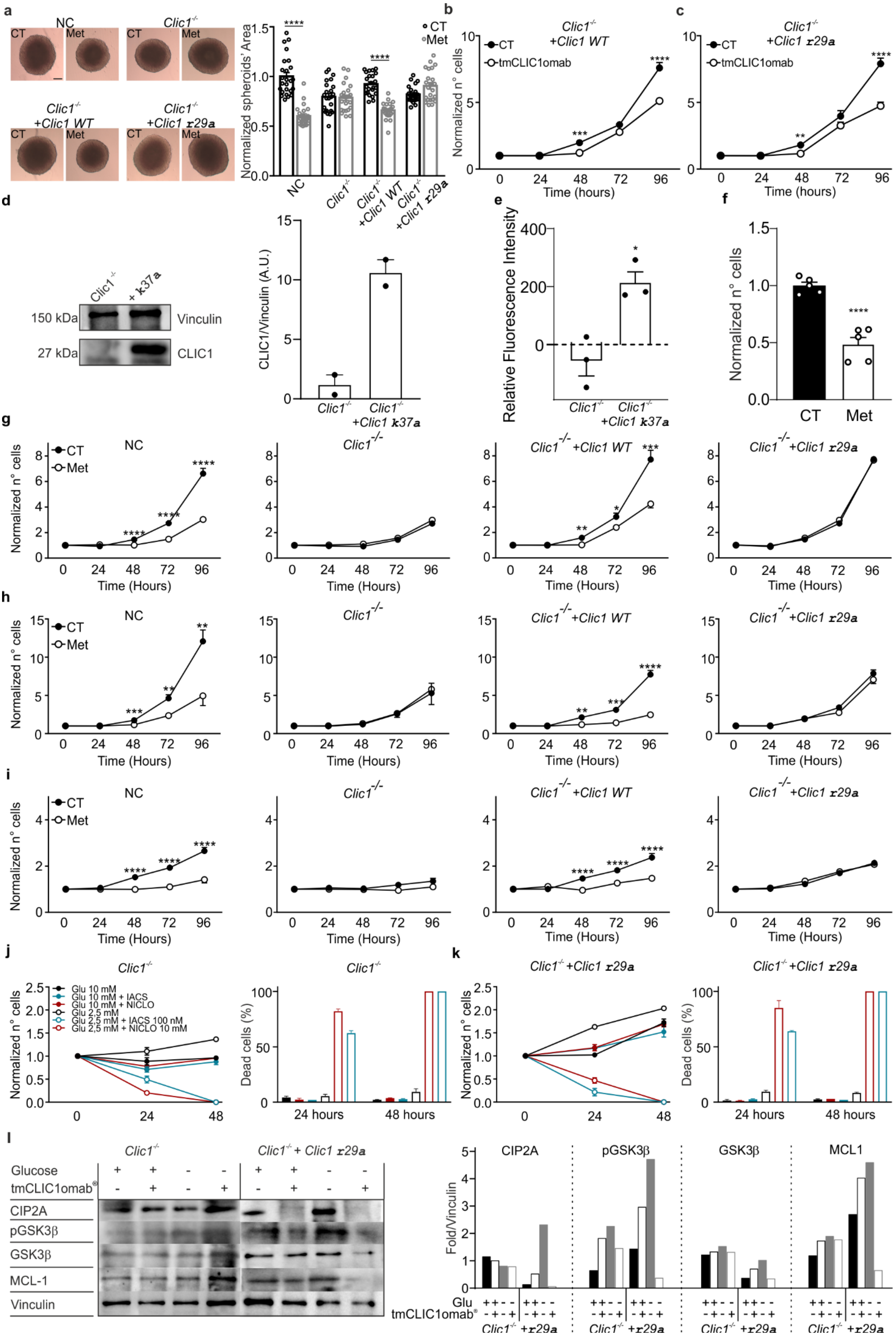
a
NC
CT
Met
Clic1-/-
Clic1-/- +Clic1 WT
Clic1-/- +Clic1 r29a
Normalized spheroids' Area
CT
Met
****
b
Clic1-/- +Clic1 WT
CT
tmCLIC1omab
Normalized n° cells
Time (hours)
c
Clic1-/- +Clic1 r29a
d
Clic1-/-
+ k37a
150 kDa
27 kDa
Vinculin
CLIC1
CLIC1/Vinculin (A.U.)
Clic1-/- +Clic1 k37a
e
Relative Fluorescence Intensity
f
Normalized n° cells
CT
Met
g
h
i
NC
Clic1-/-
Clic1-/- +Clic1 WT
Clic1-/- +Clic1 r29a
Time (Hours)
j
Glu 10 mM
Glu 10 mM + IACS
Glu 10 mM + NICLO
Glu 2,5 mM
Glu 2,5 mM + IACS 100 nM
Glu 2,5 mM + NICLO 10 mM
Dead cells (%)
24 hours
48 hours
k
l
Glucose
tmCLIC1omab®
CIP2A
pGSK3β
GSK3β
MCL-1
Vinculin
Clic1-/- + Clic1 r29a
Fold/Vinculin
CIP2A
pGSK3β
GSK3β
MCL1
Glu
tmCLIC1omab®
Clic1-/- +r29a

**Extended Data Figure 2 | CLIC1 is responsible for the impairment caused by metformin.**
**(a):** (left panel) Representative pictures showing the development of the tumor as a 3D structure (spheroid) after 96 hours of incubation in the absence or presence of 5 mM metformin as indicated in each square (right panel). Quantification of spheroids' area in the absence (black circles) or presence (grey circles) of 5 mM metformin treatment. NC: CT n=25, Met n=23; ****P<0.0001; *Clic1*$^{-/-}$: CT n=25, Met n=23*;* *Clic1*$^{-/-}$+*Clic1* WT: CT n=23, Met n=23; ****P<0.0001; *Clic1*$^{-/-}$+*Clic1* R29A: CT n=23, Met n=23; mean ± SEM, one-way ANOVA, Tukey's multiple comparison test. Scale bar 100 µm. **(b):** Growth curves of GBM3 GSCs over 96 hours in the absence (black circles) or presence of 3,5 µg/ml tmCLIC1omab antibody (empty circles) in *Clic1*$^{-/-}$+*Clic1* WT cells. CT n=8; tmCLIC1omab n=5; 48h ***P=0.0003; 96h ****P<0.0001. **(c)**: Growth curves of GBM3 GSCs over 96 hours in the absence (black circles) or presence of 3,5 µg/ml tmCLIC1omab antibody (empty circles) in *Clic1*$^{-/-}$+*Clic1* R29A cells. CT n=8; tmCLIC1omab n=5; 48h **P=0.0058; 96h ****P<0.0001. **(d)**: (left) Representative Western Blot analyses and relative quantification (right) of CLIC1 protein in K37A rescued GSCs populations' lysates.**(e):** Relative fluorescence intensity of tmCLIC1; mean ± SEM, unpaired t-test; *P=0,0127. **(f):** number of cells after 96 hours in the absence or presence of 5 mM metformin treatment in K37A rescued GSCs; mean ± SEM, unpaired t-test; ****P<0,0001. **(g-i):** Growth curves of GBM2 (g), GBM3 (h), and GL261 (i) cells over 96 hours in the absence (black circles) or presence (empty circles) of 5 mM metformin treatment in NC, *Clic1*$^{-/-}$, *Clic1*$^{-/-}$+*Clic1* WT, and *Clic1*$^{-/-}$+*Clic1* R29A, mean ± SEM, unpaired t-test. (g) NC (CT: 24,48,96h n= 12, 72h n=11; Met: 24-48h n=9, 72-96h n=8, ****P<0.0001), *Clic1*$^{-/-}$(CT: 24, 96h n=11, 48-72h n=9; Met: 24, 72, 96h n=7, 48h n=8), *Clic1*$^{-/-}$+*Clic1* WT (CT: 24-96h n=10; Met: 24, 72, 96h n=10, 48h n=9, 48h **P=0.0015, 72h *P=0,0185, 96h ***P=0,0003), *Clic1*$^{-/-}$+*Clic1* R29A (n=6). (h) NC (48h ***P=0,0005, 72h **P=0,0020, 96h **P=0,0043) *Clic1*$^{-/-}$, and *Clic1*$^{-/-}$+*Clic1* R29A (n=6), *Clic1*$^{-/-}$+*Clic1* WT (CT n=6; Met: 24-72h n=6, 96h n=5, 48h **P= 0,0021, 72h ***P=0,0002, 96h ****P<0.0001). (i) NC (CT: 24h n=9, 48-96h n=15; Met: 24h n=9, 48, 96h n=15, 72h n=14), *Clic1*$^{-/-}$(CT and Met: 24h n=6, 47-96h n=9), *Clic1*$^{-/-}$+*Clic1* WT and *Clic1*$^{-/-}$+*Clic1* R29A (n=12) ****P<0.0001. **(j):** (left) Growth curves of GBM1 GSCs over 48 hours in high (filled) and low (empty) concentration of glucose and in the absence or presence of 100 nM IACS (light blue) and 10mM niclosamide (dark red) in *Clic1*$^{-/-}$ (n=3, 48h IACS and niclosamide ****P<0.0001); mean ± SEM, one-way ANOVA, Tukey's multiple comparison test. (right) Analysis of the percentage of dead cells shown in left panel (n=3; 24-48h IACS and niclosamide ****P<0.0001); mean ± SEM, one-way ANOVA, Tukey's multiple comparison test. **(k):** (left) Growth curves of GBM1 GSCs over 48 hours in high (filled) and low (empty) concentration of glucose and in the absence or presence of 100 nM IACS (light blue) and 10mM niclosamide (dark red) in *Clic1*$^{-/-}$+*Clic1* R29A (n=3, 48h IACS and niclosamide ****P<0.0001); mean ± SEM, one-way ANOVA, Tukey's multiple comparison test. (right) Analysis of the percentage of dead cells shown in left panel (n=3; 24-48h IACS and niclosamide ****P<0.0001); mean ± SEM, one-way ANOVA, Tukey's multiple comparison test. **(l):** Representative Western blot analysis (left) and its quantification (right) of *Clic1*$^{-/-}$ and *Clic1*$^{-/-}$ +Clic1 R29A cultured in high (black filled column) and low (grey filled column) concentration of glucose in the absence (black empty column) or presence (grey empty column) of 3,5 µg/ml of tmCLIC1omab®.

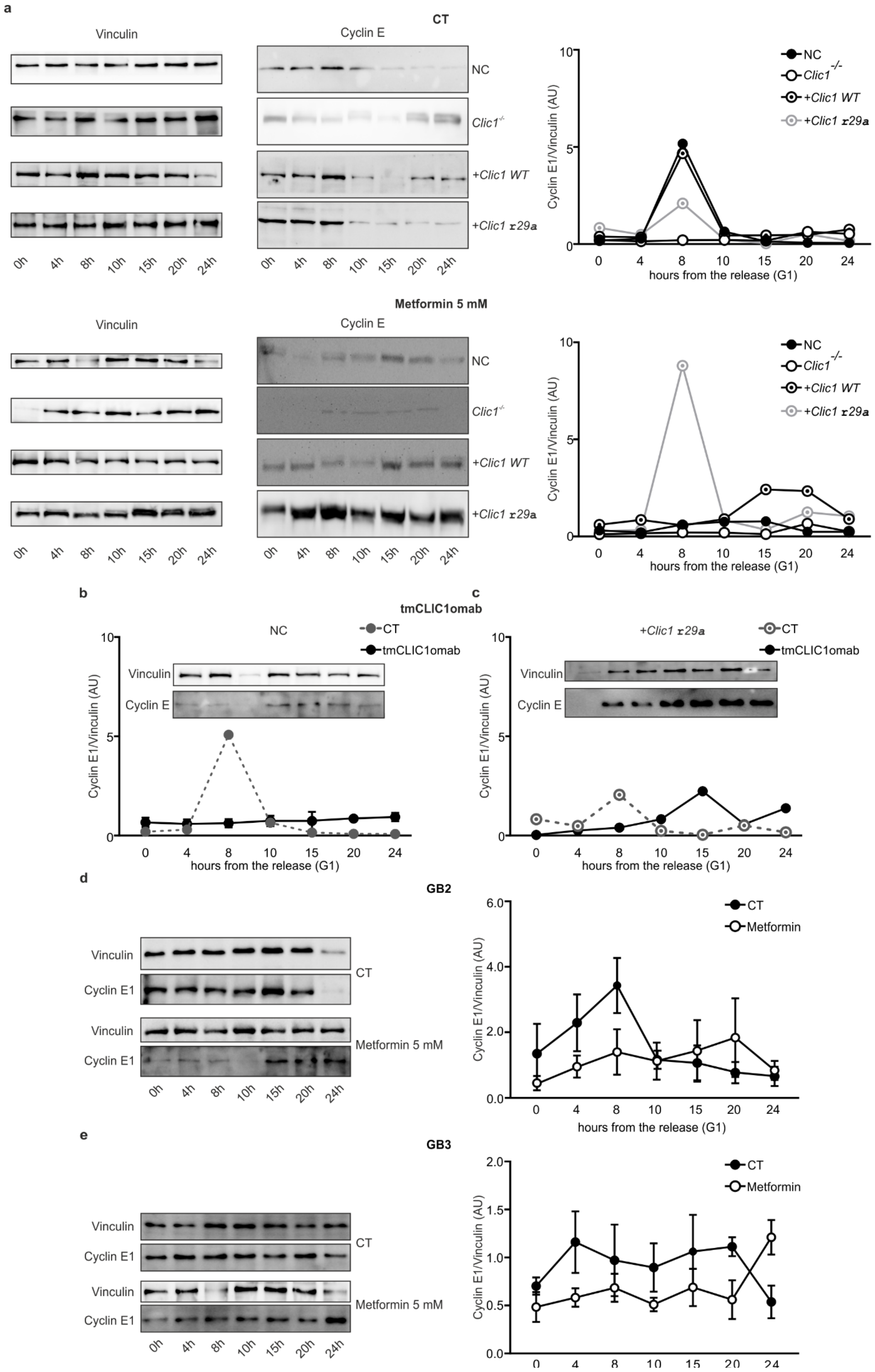
a
CT
Vinculin
Cyclin E
NC
Clic1-/-
+Clic1 WT
+Clic1 r29a
0h 4h 8h 10h 15h 20h 24h
Cyclin E1/Vinculin (AU)
hours from the release (G1)
Metformin 5 mM
b
tmCLIC1omab
NC
CT
tmCLIC1omab
Vinculin
Cyclin E
c
+Clic1 r29a
d
GB2
Cyclin E1
Metformin
e
GB3

**Extended Data Figure 3 | Cyclin E expression in GBM1, GBM2, and GBM3 primary cultures.**
**(a):** (left) Cyclin E1 expression representative Western Blot analyses of lysates of NC, *Clic1*$^{-/-}$, and rescued GBMM1 GSCs populations at different time points after the release from G1 synchronization in absence (top) or presence (bottom) of 5mM metformin treatment. In the right panel is plotted the trend of cyclin E1 protein expression in the samples (n=3).
**(b):** (top) Cyclin E1 expression representative Western Blot analyses of lysates of NC (left) and *Clic1*$^{-/-}$+*Clic1* R29A (right) GBM1 populations at different time points after the release from G1 synchronization in presence of 3,5 µg/ml tmCLIC1omab (bottom) trend of cyclin E1 protein expression in the treated samples (filled circles) (n=3). Dashed lines refer to untreated samples shown in (a).
**(c):** (left) Cyclin E1 expression representative Western Blot analyses of lysates of NC GBM2 (top) and GBM3 (bottom) populations at different time points after the release from G1 synchronization in absence or presence of 5mM metformin treatment. In the right panel is plotted the trend of cyclin E1 protein expression (n=3) in control (black circles) and metformin-treated cells (empty circles).

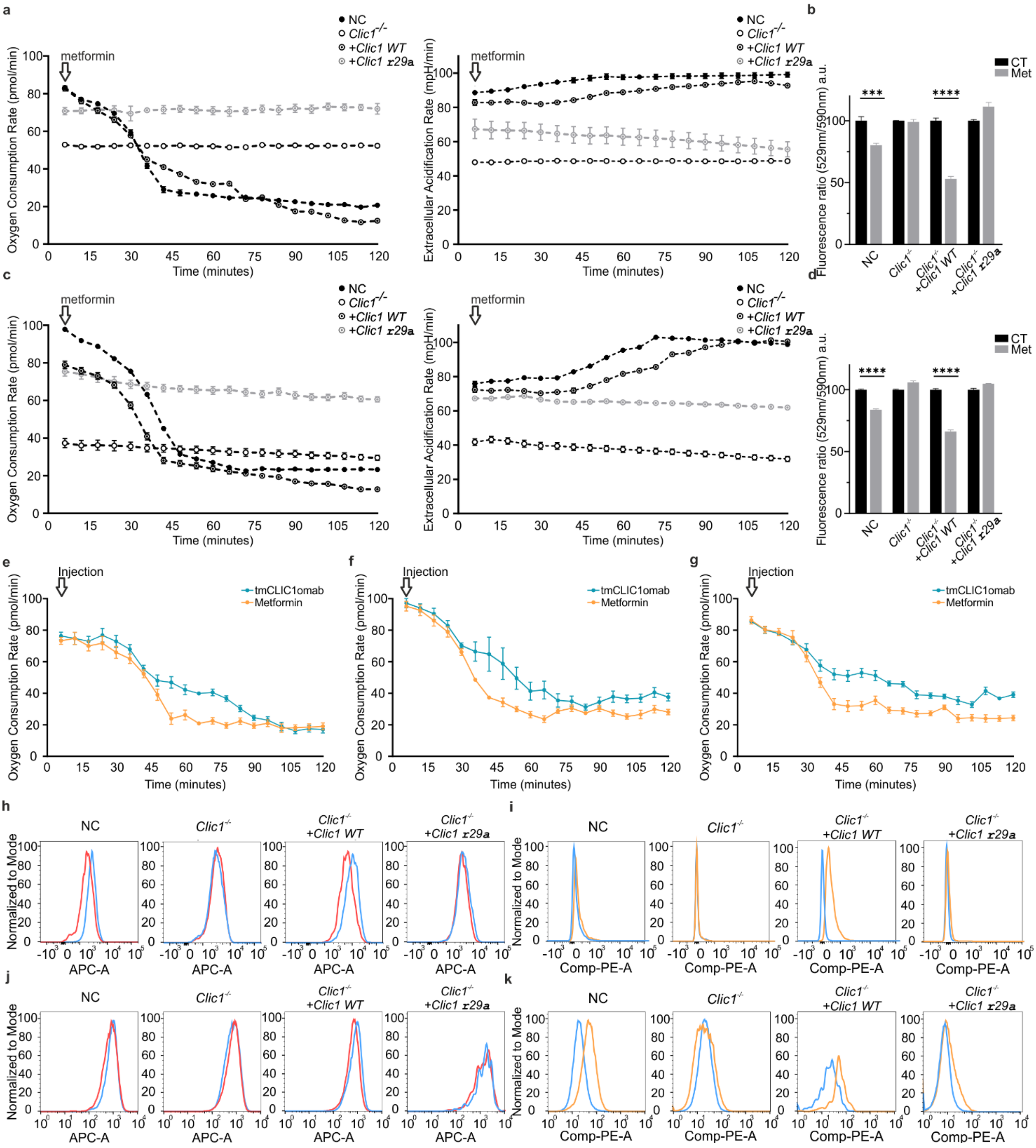


**Extended Data Figure 4 | Metformin's effect on metabolism depends on CLIC1**

**(a) and (c):** Oxygen consumption rate (left) and Extracellular acidification rate (right) in GBM2 (a) and GBM3 (c) as an indicator of oxidative phosphorylation and glycolytic activity after acute injection of 10 mM metformin in NC (black circles), *Clic1*$^{-/-}$ (empty circles), *Clic1*$^{-/-}$ +Clic1 WT (black circle dot), *Clic1*$^{-/-}$ +Clic1 R29A cells (gray circle dot). Each experimental point was taken every 6 minutes. **(b and d):** Mitochondrial membrane potential measured in GBM2 (b) and GBM3 (d) through JC-1 probe in NC, *Clic1*$^{-/-}$, *Clic1*$^{-/-}$ +Clic1 WT, and *Clic1*$^{-/-}$ +Clic1 R29A in the absence (black columns) or presence (grey columns) of 5 mM metformin treatment. n=2; GBM2, NC: CT vs Met, **P=0.0008; *Clic1*$^{-/-}$+*Clic1* WT: CT vs Met, ****P<0.0001; GBM3, ****P<0.0001 mean ± SEM, two-way ANOVA, Sidak's multiple comparison test. **(e-g):** Oxygen consumption rate in GBM1 (e), GBM2 (f), and GBM3 (g) after acute injection of 10 mM metformin (orange) and 3,5 µg/ml tmCLIC1omab antibody (cyan). **(h and j):** Representative plot of FACS analysis of

CellROX™ Deep Red Reagent fluorescence for oxidative stress detection in GBM2 (h) and GBM1 (j). Cells were incubated for 3 hours in the absence (blue) or presence (red) of 5mM of Metformin. i and k. Representative plot of FACS analysis of MitoSOX™ Red Indicator fluorescence for Mitochondrial Superoxide detection in GBM2 (i) and GBM1 (k). Cells were incubated for 3 hours in the absence (blue) or presence (orange) of 5mM of Metformin.

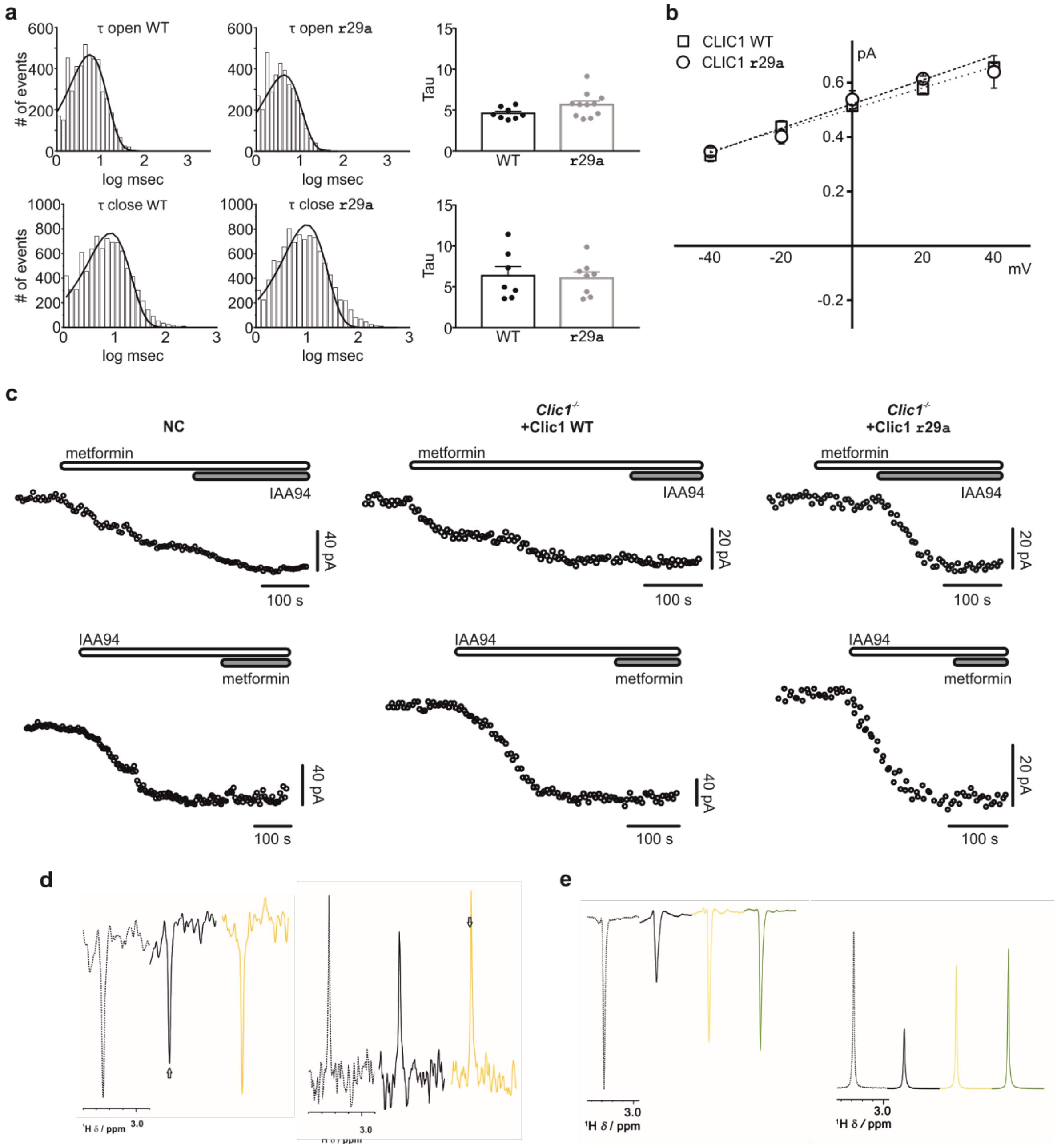


**Extended Data Figure 5 | Metformin directly interacts with tmCLIC1.**

**a:** (left) Open and Close times (t) of tmCLIC1 channel in *Clic1*$^{-/-}$+*Clic1* WT and *Clic1*$^{-/-}$+*Clic1* R29A GBM3 cells measured during at least 3 minutes of continue recordings in outside-out configuration. Histograms were best fitted by a single exponential decay function (bold line). (right) Quantification of single Tau values. t open *Clic1*$^{-/-}$+*Clic1* WT 4.59 ± 0.22 ms (n=9); *Clic1*$^{-/-}$+*Clic1* R29A 5.65 ± 0.46 ms (n=11). t close *Clic1*$^{-/-}$+*Clic1* WT 6.35 ± 1.13 ms (n=7); *Clic1*$^{-/-}$+*Clic1* R29A 6.06 ± 0.75 ms (n=8). (Open: *Clic1*$^{-/-}$+*Clic1* WT n=8 *Clic1*$^{-/-}$+*Clic1* R29A n=11; Close: *Clic1*$^{-/-}$+*Clic1* WT n=7 *Clic1*$^{-/-}$+*Clic1* R29A n=8).

**b:** Current-voltage relationship of tmCLIC1 single channel outside-out experiments. The calculated conductance was 3.89 ± 0.21 pS for *Clic1*$^{-/-}$+*Clic1* WT and 4.46 ± 0.27 pS for *Clic1*$^{-/-}$+*Clic1* R29A.
**c:** Representative time-course of whole cell currents in NC and WT/R29A rescued cells. Each point represents the average current of the last 100 ms of a single current trace. Cells were stimulated every 5 seconds with a 800 ms, +60 mV test potential from the resting potential. Once the current amplitude reached a constant value, metformin 5mM and IAA94 100μM (top) and vice versa (bottom) were perfused.
**d:** WaterLOGSY, e. $T_2$ filter $1^H$ NMR spectra of 50 μM metformin in absence (dashed line) and in presence of 10 μM CLIC1 wild type (black) R29A mutant (yellow). Only the spectral region containing the metformin dimethyl resonance signal are displayed superimposed and shifted. Arrows highlight signal changes showing binding of metformin to CLIC1 wild type; no changes are observed in the presence of the R29A mutant.
**e:** same procedure as Figure 3b but with twice cell number.
